# Harnessing Emergent Properties of Microbial Consortia: Assembly of the Xilonen SynCom

**DOI:** 10.1101/2024.04.24.590952

**Authors:** Gabriela Gastélum, Bruno Gómez-Gil, Gabriela Olmedo-Álvarez, Jorge Rocha

**Affiliations:** Centro de Investigación en Alimentación y Desarrollo A.C. (CIAD) Unidad Regional Hidalgo. San Agustín Tlaxiaca, Hidalgo, 42162, Mexico; Centro de Investigación en Alimentación y Desarrollo A.C. (CIAD) Unidad Mazatlán. Acuicultura y Manejo Ambiental. Mazatlán AP711, Sinaloa, Mexico; Departamento de Ingeniería Genética, CINVESTAV-Unidad Irapuato, Guanajuato, Mexico; Programa de Agricultura en Zonas Áridas, Centro de Investigaciones Biológicas del Noroeste, La Paz, Baja California Sur, 23096, Mexico

**Keywords:** synthetic communities, SynCom, seed-endophytes, inoculants, emergent properties, biofilm formation, native maize landraces

## Abstract

Synthetic communities (SynComs) are valuable tools for addressing fundamental questions in microbial ecology regarding community assembly. They could also potentially aid in successfully manipulating microbial communities for clinical, biotechnological, and agricultural applications. SynCom design is complicated since interactions between microbes cannot be predicted based on their individual properties. Here, we aimed to assemble a higher-order SynCom from seed-endophytic bacteria isolated from native maize landraces. We screened co-cultures that included strains from the Bacilli class, and the *Burkholderia* and *Pseudomonas* genera since these taxa have been previously shown to be important for the fertility of native maize landraces. We developed a combinatorial, bottom-up strategy aimed at the detection of a complex colony architecture as an emergent collective property. Using this simplified approach, we assembled a SynCom composed of *Bacillus pumilus* NME155, *Burkholderia contaminans* XM7 and *Pseudomonas* sp. GW6. The strains exhibited positive and negative interactions when evaluated in pairs, but their higher-order assembly results in a complex colony architecture, which is considered a proxy of biofilm formation. This SynCom was named *Xilonen* after the Aztec goddess of young maize and fertility. The *Xilonen* SynCom will aid in studying the molecular and ecological basis mediating maize fertility.

## Introduction

Mesoamerica holds special significance in the history of agriculture as the place where plants such as maize, beans, chili peppers, and squash – now staple foods for the population – were domesticated. Mexico still maintains dozens of native maize landraces and preserves the agricultural practice of cultivating them in a polyculture system known as *milpa* (1, 2). *Milpa* productivity is achieved without the use of pesticides, fertilizers, irrigation, or mechanization (3), and traditional agronomic practices are central not only to nutrition and economy but also to culture, traditions, and the general worldview of indigenous communities in Mexico (4). Indeed, crop success was attributed to deities such as Xilonen, the Aztec goddess of young maize and fertility (5), and still, understanding the ecological basis of plant fertility in *milpas* is highly relevant to develop novel strategies for sustainable agriculture.

Microbes are essential for plant health and crop productivity since they can aid the plant in nutrient acquisition (6), biotic and abiotic stress alleviation (7), and immunomodulation (8). Soil microbes are attracted to root exudates (9), which can sustain the growth of specific bacterial groups that perform these functions (10, 11). Root exudates can also induce biofilm formation (12), a process that is also affected by interactions with other microbes (13–15) and is indispensable for root colonization and the establishment of plant-microbe interactions.

Our recent studies in native maize from *milpa* traditional agroecosystems revealed that seed-endophytic bacteria that remain on the seedling root upon germination appear to be particularly important for the fertility of these plants (16–18). In this fraction of the bacteriome, Bacilli strains are highly diverse and abundant, and are capable of colonizing the seedling root (16). Also, seed-endophytic *Burkholderia* spp. can be highly antagonistic and could mediate community assembly and biocontrol (17). Finally, *Pseudomonas* spp., contribute to drought tolerance in landraces from arid *milpas* (18). Since the functions attributed to these bacterial genera were described for axenic cultures, it is still necessary to generate more complex experimental systems that allow 1) grasping community-level functions and 2) understanding the basis of community assembly. This information will aid in the study of interactions that mediate maize fertility and could be useful resources for sustainable agriculture.

The use of microbial products in agriculture is a promising approach to aid plant health while reducing the use of chemical pesticides and fertilizers (19). Notably, the use of microbial consortia allowed increased performance in laboratory settings, compared to single strains (20). However, scaling to field applications is still a limiting step (21–24) probably due to the deficient incorporation of microbial ecology approaches in the development of multistrain products (25, 26). In the formulation of microbial consortia, the typical approach involves mixing ‘biocompatible’ strains, which entails selecting microbes based on complementary functions identified through the study of strains in isolation, and including only those that do not demonstrate antagonism when paired together (27–36). This method lacks a solid foundation, since antagonism tends to be attenuated in higher-order systems (37, 38). Also, the functioning of the plant microbiota results from complex interactions including metabolic cross-feeding, competition and growth inhibition between community members (39), all of which are unpredictable from the characterization of individual strains (39–41). Novel approaches in the design of microbial consortia for agriculture should consider the ecological processes governing community assembly and stability, and their impact on the resulting beneficial functions.

Synthetic communities (SynComs) have emerged as valuable tools for establishing causal relationships between members (or their genes) and community functions (phenotypes) (42, 43). These laboratory-assembled communities can be designed to represent natural communities but with reduced complexity (42), thereby enabling precise experimental manipulation aimed at generating microbial consortia for enhancing plant health and productivity (44–46). However, the design of simplified, tractable and ecologically relevant SynComs is still challenging (39–41). Bottom-up approaches for SynCom design represent a rational strategy where isolates are used as building blocks (42, 43). However, most strategies use technically challenging co-culturing and analysis strategies (47–50) that are inaccessible for many non-specialized laboratories working with natural isolates.

Recently, bacterial co-culture studies have shown that collective functions can emerge due to strain-specific interactions (13, 37, 38, 51, 52). Collective functions can influence the fitness of bacterial populations in the rhizosphere impacting host health (53–56) and their macroscopic manifestations can be easily detected *in vitro*. Importantly, emergent collective functions in a co-culture indicate strain co-existence, community assembly, and the presence of bacterial interactions that result in a beneficial function. However, these traits have not been widely exploited for SynCom assembly (57).

Here we aimed to generate a SynCom using seed-endophytic bacteria from native maize, as a model to study the beneficial functions of bacteria from *milpas*. To achieve this, we devised a simplified combinatorial bottom-up approach, followed by the identification of collective emergent functions within co-cultures. We focused on emergent complex architecture in mixed colony biofilms, a trait that could be relevant for root colonization. Our screening resulted in the assembly of a three-member community named Xilonen (5). Pairwise interactions and community dynamics were evaluated as a first step for establishing this community as a model. Overall, the Xilonen community is a valuable tool to grasp the molecular and ecological basis mediating the contribution of the microbiota to native maize fertility. Moreover, this work provides a framework for rapid, design-free, accessible assembly of SynComs with ecological, biotechnological, and clinical relevance.

## Results

### 1. Assembly of the Xilonen synthetic community

To develop synthetic communities (SynComs), we established a bottom-up combinatorial strategy based on the detection of emergent properties in bacterial co-cultures. Our screening focused on detecting emergent complex architecture on colony biofilms since these macroscopic structures are associated with the synthesis of molecules mediating biofilm formation and other multicellular behaviors of bacteria (reviewed in refs 54 and 55).

We used 27 seed-endophytic bacterial strains from maize landraces from our collection, including 20 *Bacillus* sp., 3 *Burkholderia* sp., and 4 *Pseudomonas* sp. strains (Supplementary Table S1). These bacterial taxonomic groups have been previously found to be relevant for the fertility of native maize varieties (16–18). It is important to note that a bottom-up approach testing all combinations of size *k* = {2, 3, 4} from 27 strains would entail preparing and screening 20,826 co-cultures. We reduced this number by 1) testing only co-cultures with at least two bacterial genera, and 2) testing pairwise interactions first, selecting specific communities of *k* = 2 with emergent colony morphology before introducing an additional strain to assemble communities of *k* = 3, and screening for an emergent morphology differing from that of the pairwise co-cultures (Fig. 1a and 1b).

**Figure 1.**
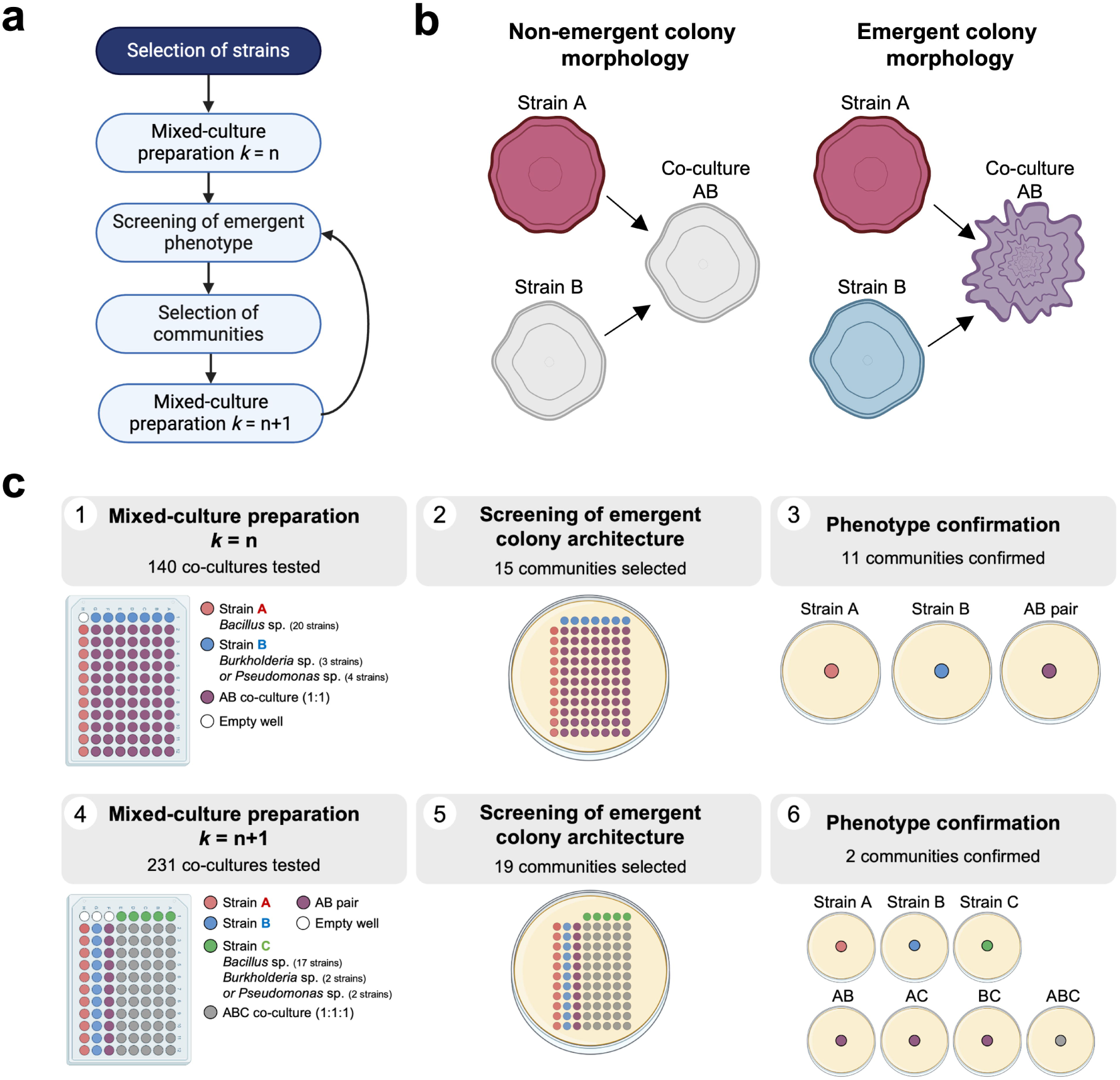
Assembly of the Xilonen synthetic community through screening for emergent colony architecture resulting from interactions between seed-endophytic bacteria of native maize landraces. **a,** The general strategy was based on preparation of pairwise co-cultures and screening of emergent functions for the assembly of synthetic bacterial communities of size (*k*) and subsequent addition of members for *k*+1 communities and screening. **b,** Representation of emergent colony morphology considered in our screening.

Non-emergent colony morphology: morphology in the co-culture of two strains (AB) is similar to morphology of strain A or B (left). Emergent colony morphology: morphology in the co-culture AB differs from morphologies A and B (right). **c,** Workflow followed for the assembly of the Xilonen synthetic community. First, AB pairs were mixed using two bacterial genera (1). After incubation, colonies were screened for emergent colony morphology (2), and phenotypes were confirmed in separate plates (3). Then, a third strain C was added to selected AB pairs (4), and co-cultures *k* = 3 were screened for emergent colony morphology (5). Finally, phenotypes of *k* = 3 communities were confirmed in separate plates against all possible single and pairwise co-cultures (6). Created with BioRender.com.

First, we prepared communities of *k* = 2 including one of 20 *Bacillus* sp. (strain A) and one of seven *Burkholderia* sp. or *Pseudomonas* sp. (strain B) (Fig. 1c-1, Supplementary Fig. S1a). From the 140 co-cultures, we detected 15 AB pairs exhibiting emergent colony morphology (Fig. 1c-2). When these communities were tested in separate agar plates, 11 communities with emergent morphology were confirmed (Fig. 1c-3). Representative colony biofilms of communities *k* = 2 with emergent morphology are shown in Supplementary Fig. S1b.

**Figure 2.**
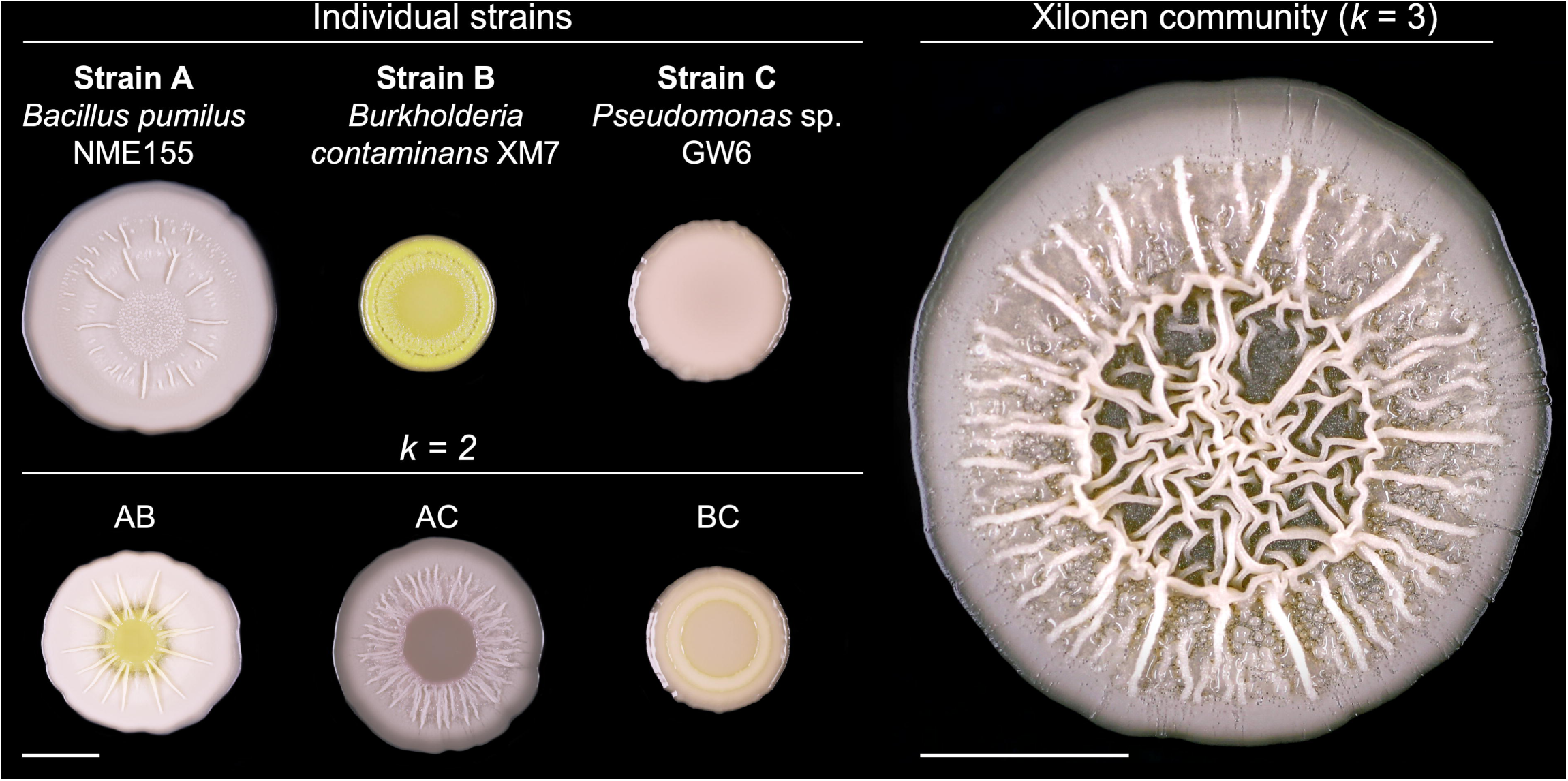
Xilonen synthetic community. Complex colony architecture emerges only through the interaction of the three strains in the community. Colonies of individual strains that form the synthetic community, colonies of pairwise co-cultures (*k* = 2), and the Xilonen community (*k* = 3). Each colony was grown individually in separate LB plates. Pictures were taken after 3 days of incubation at 30 °C. Both scale bars are 5 mm.

**Figure 3.**
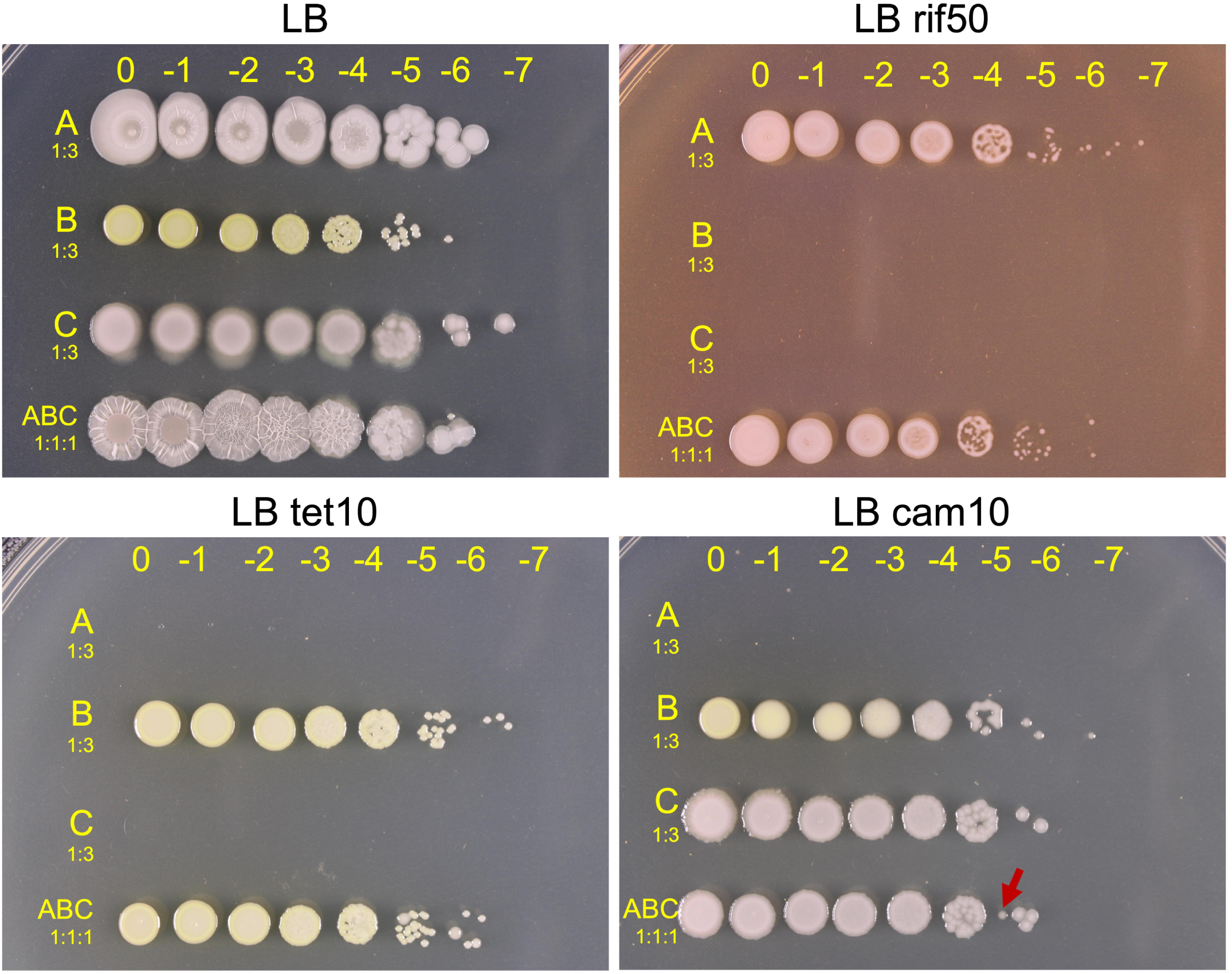
Selective media for individual CFU counts of each member of the community. Spot inoculations of 10-fold serial dilutions of liquid cultures of individual strains and mixed cultures. Overnight bacterial liquid cultures were washed with PBS and the optical density at 600 nm was adjusted to 1 ± 0.1. Washed bacterial cultures were mixed at a 1:1:1 volume ratio, while individual strains were diluted 1:3 in sterile PBS. Five μL of 10-fold serial dilutions were spotted on LB and each selective medium. Rifampicin 50 μg/mL (rif50); tetracycline 10 μg/mL (tet10) or chloramphenicol 10 μg/mL (cam10) were added to LB medium to recover strains A (Bp_NME155), B (Bc_XM13), and C (P_GW6) respectively. The arrow in the LB cam10 plate indicates small colonies corresponding to strain B, which are not considered for CFU counts in this selective medium.

Next, we screened 231 co-cultures of *k* = 3 that resulted from adding 21 strains (strain C) to each of the 11 confirmed AB communities (Fig. 1c-4, Supplementary Fig. S2a). In this step, we identified 19 ABC co-cultures with emergent colony morphology (Fig. 1c-5). When confirming their phenotype in separate plates, we found that only two ABC communities exhibited a robust emergent colony morphology. Both morphologies differed from individual strains (A, B, C) and from the pairwise co-cultures (AB, AC, and BC, Fig. 1c-6); therefore, they were considered as emergent morphologies resulting from higher-order interactions, *i.e*., those that arise in communities of three or more strains, and induce emergent properties not observed in the individual strains (60). One of the two communities showed emergent spreading; it consisted of *Bacillus* sp. NME63, *Pseudomonas* sp. GW1, and *Bacillus* sp. NME246 (Supplementary Fig. S2b). The other community exhibited emergent colony architecture (Fig. 2). This community included the strains *Bacillus* sp. NME155, *Burkholderia* sp. XM7 and *Pseudomonas* sp. GW6 (Fig. 2). We also screened 24 co-cultures of *k* = 4 by adding 12 strains to both communities of *k* = 3, but we did not observe an emergent morphology in any of the co-cultures (not shown).

**Figure 4.**
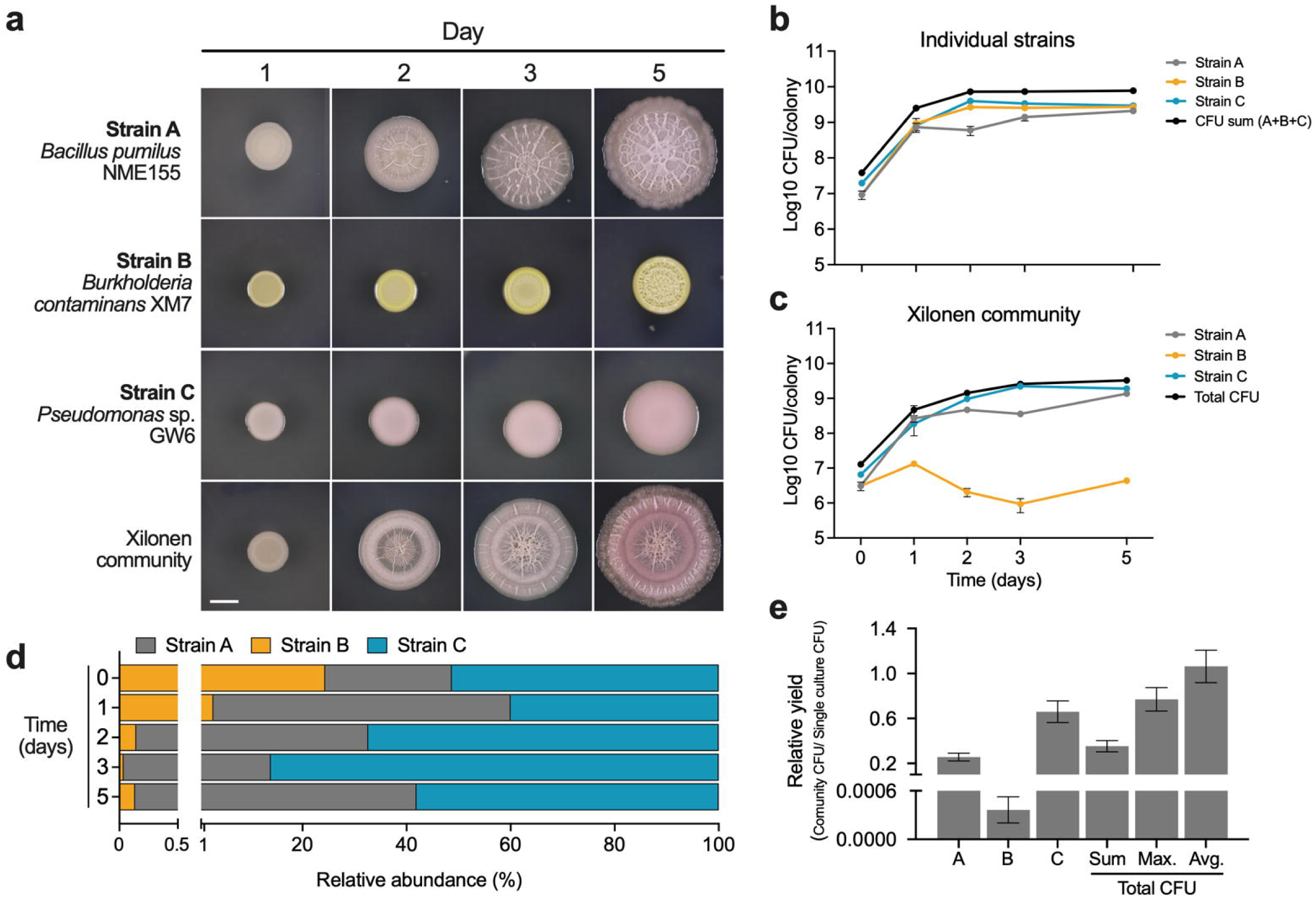
Growth dynamics of individual strains in the Xilonen SynCom. **a,** Colony development of individual strains and the mixed community. Scale bar, 5 mm. **b,** CFU counts from colonies of individual strains. **c,** CFU counts of individual strains in the Xilonen community. **d,** Relative abundance of each member of the community on days 0, 1, 2, 3, and 5. **e,** Relative yield of individual strains and total cells in the Xilonen community, compared to individually grown strains on day 3. Total CFUs in (e) are shown relative to the sum of individual strains (Sum), to the strain with maximum individual yield (Max.) and the average yield per colony (Avg.). The average of three replicates ± SD is shown.

**Figure 5.**
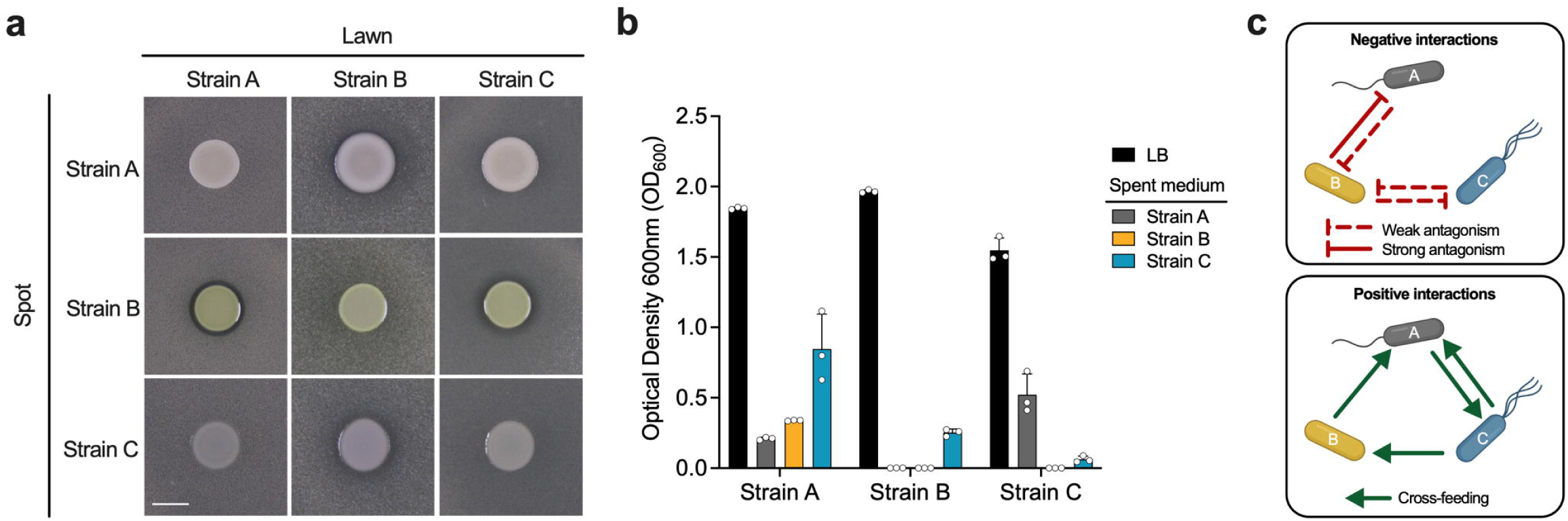
Pairwise negative (antagonism) and positive (cross-feeding) interactions between strains in the synthetic community. **a,** Antagonistic interactions using the spot-on-lawn assay. Pictures were taken 24 h after incubation at 30 °C. Scale: 5 mm. **b,** Cross-feeding interactions were evaluated by measuring bacterial growth in cell-free spent medium. Each strain was grown in fresh LB and cell-free spent medium (see methods). After 2 days of incubation at 30 °C and 200 rpm, OD_600_ was measured. Dots represent individual replicates. **c,** Schematic representation of pairwise antagonistic (top) and cross-feeding interactions (bottom). Strain A: *Bacillus pumilus* NME155; Strain B: *Burkholderia contaminans* XM7; Strain C: *Pseudomonas* sp. GW6.

**Figure 6.**
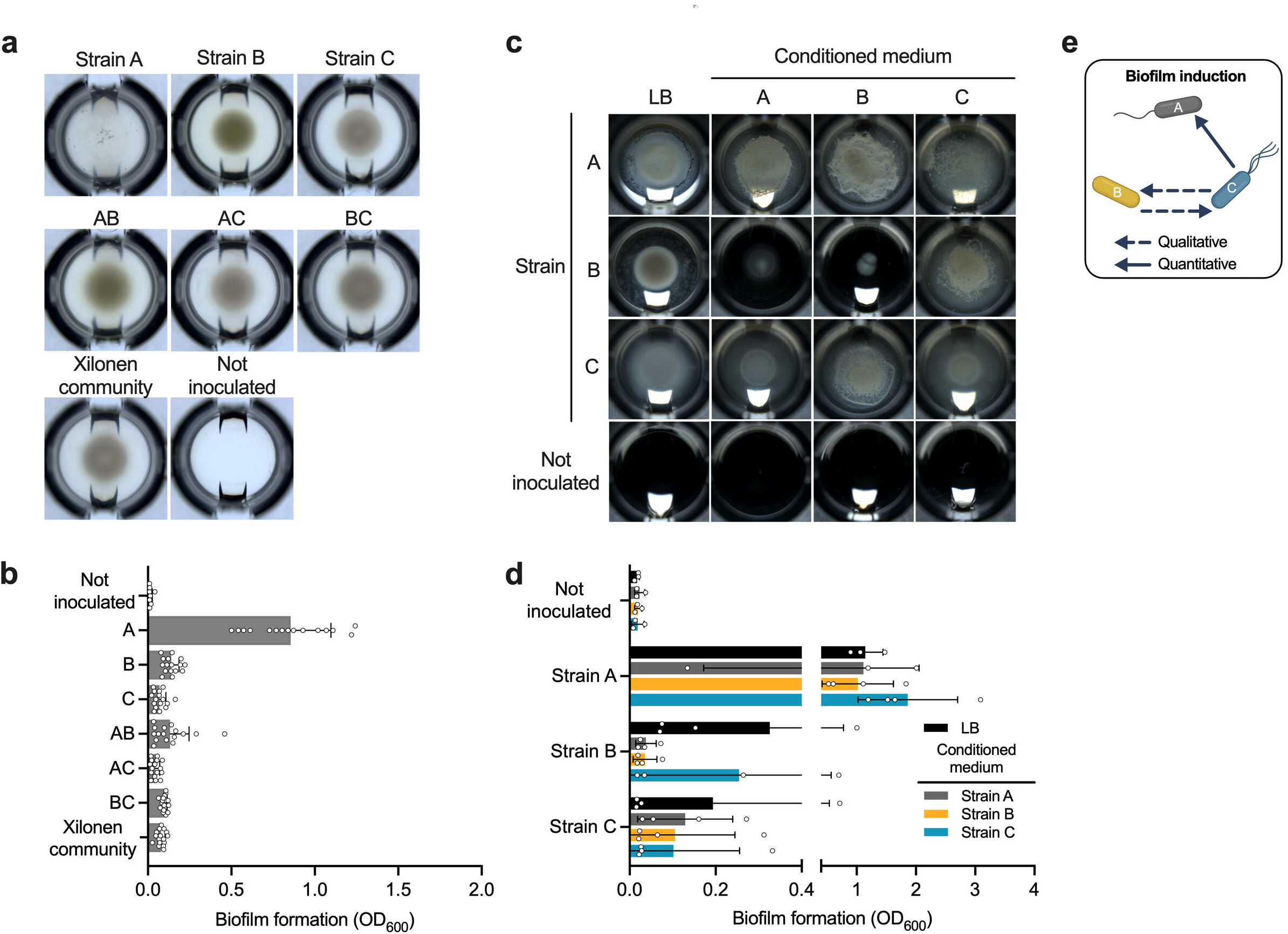
Pairwise induction of biofilm formation in the community in standing liquid culture. **a,** Microtiter wells showing the growth of single strains, pairwise co-cultures, and the three-member SynCom in standing liquid LB medium. Plates were imaged after 3 days of incubation at 30 °C. **b,** Biofilm formation of cultures shown in (a) was quantified using the crystal violet method. **c,** Microtiter wells showing the development of pellicles of each strain grown individually in fresh LB medium and in conditioned medium (see methods). Plates were imaged after 3 days of incubation at 30 °C. **d,** Biofilm formation of cultures shown in (c) was quantified using the crystal violet method. **e,** Schematic representation of pairwise induction of biofilm formation. Strain A: *Bacillus pumilus* NME155; Strain B: *Burkholderia contaminans* XM7 and Strain C: *Pseudomonas* sp. GW6. Dots represent individual replicates.

In summary, our exploration of interactions and colony architecture in communities of seed-endophytic bacteria from native maize included the screening of 395 co-cultures, resulting in the identification of one community *k* = 3 that exhibited an emergent complex colony architecture and included the three bacterial taxa considered (Fig. 2). We named this community Xilonen, and it was further characterized.

### 2. Genomic sequence and species-level identification of the members of the Xilonen community

The Xilonen synthetic community is composed of *Bacillus* sp. NME155, *Burkholderia* sp. XM7, and *Pseudomonas* sp. GW6 strains, which were preliminarily identified at the genus level in previous studies (16–18). For species-level identification, we carried out genomic DNA sequencing and *de novo* assembly. Sequencing, assembly, and annotation statistics are summarized in Supplementary Table S2. From the genomic data, two of the strains were identified at the species level as *Bacillus pumilus* (Bp_NME155) and *Burkholderia contaminans* (Bc_XM7). Their Average Nucleotide Identity (ANI%) and digital DNA-DNA Hybridization (dDDH%) values were 95.36% and 63.9% for Bp_NME155; 99.92% and 98.9% for Bc_XM7, respectively. The *Pseudomonas* sp. GW6 strain could only be identified at the genus level since it had an ANI of only 91.89% and 44.8% dDDH when compared with its closest relative *Pseudomonas sediminis* PI11. These values are below the threshold recommended for species identification (≥ 95% ANI and ≥ 70% dDDH) (61, 62) and thus, although further analyses will be needed, *Pseudomonas* sp. GW6 (P_GW6) could be considered a new species.

Results from the quality evaluation of the assemblies showed that Bp_NME155 and P_GW6 (GenBank Acc. No. SAMN39859859 and SAMN39859861) had 100% completeness with 0.03% and 0.38% contamination, respectively (Supplementary Table S2). For Bp_NME155 4,056 coding sequences, 86 RNAs, and 1,385 annotated genes were predicted; P_GW6 exhibited 5,489 coding sequences, 79 RNAs, and 1,591 annotated genes (Supplementary Table S2). The assembly of Bc_XM7 displayed a lower quality of 66.39% completeness and 7.48% contamination, hence gene annotation and GenBank submission were not possible. Lower quality in the assembly of Bc_XM7 could be explained by the presence of two tightly associated and taxonomically close strains in the original XM7 isolate. This bacterial culture exhibits two colony morphologies consisting of small and large colonies (Supplementary Fig. S3a) which could not be independently cultured after subsequent plating and selection (Supplementary Fig. S3b). We expect that sequencing with longer reads could improve the quality of sequences for further genomic characterization of the XM7 strain or consortium (63, 64).

### 3. Growth dynamics in the Xilonen community

To examine the growth dynamics in the community, we devised a strategy to recover and quantify each strain from the mixed colony biofilm. Differential antibiotic resistance was deemed a useful approach. First, we performed Minimum Inhibitory Concentration (MIC) assays using eight antibiotics (Supplementary Fig. S4). These assays indicated that tetracycline and chloramphenicol could be used to generate selective media for Bc_XM7 and P_GW6, respectively. Next, we tested if we could quantify each strain from a mixture of independently grown liquid cultures. The addition of 10 µg/ml of tetracycline allowed the selection of Bc_XM7, completely inhibiting the growth of Bp_NME155 and P_GW6 (Fig. 3). Also, 10 µg/ml of chloramphenicol was used to recover P_GW6 while inhibiting the growth of Bp_NME155 and decreasing the size of Bc_XM7 colonies. In each case, the corresponding strain grew to similar CFUs compared to medium without antibiotics (Fig. 3).

Our MIC assays did not allow the generation of a selective medium for Bp_NME155. Hence, for this strain, we obtained a rifampicin-resistant variant (rif^R^) to allow its selection and quantification from the colony biofilm (see methods). We verified that this variant maintained the colony morphology of the WT strain, and allowed the development of emergent complex colony architecture when incorporated in the Xilonen community (Supplementary Fig. S5). The addition of 50 μg/mL of rifampicin allowed the quantification of Bp_NME155 rif^R^ from the mixture of three strains (Fig. 3).

Next, we tested whether the selective media allowed accurate CFU counts in suspensions obtained from colony biofilms of individual strains. Strains Bp_NME155 and Bc_XM7 showed consistent CFU counts in media with and without the corresponding antibiotics when measured from colony biofilms (not shown). However, the addition of chloramphenicol resulted in a decrease in CFU counts of P_GW6 when it was quantified from colony biofilms (Supplementary Fig. S6). This phenomenon likely affects the quantification of P_GW6 from the Xilonen community; however, our preliminary assays showed that P_GW6 was the dominant species in the mixed biofilm, hence, it was possible to quantify its CFUs in non-selective medium from its distinct colony morphology.

We used the devised differential antibiotic resistance method to evaluate the growth dynamics of the three strains in colony biofilms of single strain cultures compared to the community context (Fig. 4a), following CFUs during 5 days. Single-strain colony biofilms were initially inoculated with 9.33×10^6^ CFU/colony of Bp_NME155 (strain A), 9.33×10^6^ of Bc_XM7 (strain B), and 1.96 ×10^7^ of P_GW6 (strain C) In these individual inoculations, colonies of strain B and C presented a maximum of 2.66×10^9^, and 3.96×10^9^ CFU/colony on day 2, respectively; strain A presented a maximum of 2.1×10^9^ CFU/colony on day 5 (Fig. 4b).

Interactions in the community context caused changes in the growth dynamics of the strains (Fig. 4b and 4c). At day 0, the community was inoculated with 1.28×10^7^ total CFU/colony, consisting of 3.11×10^6^ CFU of strain A, 3.11×10^6^ CFU of strain B, and 6.55×10^6^ CFU of strain C (Fig. 4c). The maximum CFU count was observed on day five with a total yield of 3.39×10^9^ CFU/colony (1.36×10^9^ CFU of strain A, 4.36×10^6^ CFU of strain B and 1.9×10^9^ CFU of strain C) (Fig. 4c). In the inoculation mixture, both strains A and B represented 24.35% each, and strain C comprised 51.3% of the community. Strain C dominated the community during most days of biofilm development ranging from 40% to 86.08%. Abundance of strain A ranged from 13.87% to 57.14%, dominating the community on day two. Strain B was the least abundant in the community with 0.036% to 2.85% (Fig. 4d).

Emergent biofilm architecture was observed on day 3 (Fig. 4a), however, interactions in the community and the emergence of colony architecture were not accompanied by growth induction (65) of the community or the individual strains. On day 3, the growth of strains A, B, and C in the community represented only 26%, 0.036%, and 66.33%, respectively, compared to their growth after three days as individual colonies (Fig. 4e). The total yield in the Xilonen community on day 3 (2.59×10^9^ CFU/colony) was only 35% of the expected yield from the sum of the three strains grown individually, 77% of that of the strain with greater individual yield (strain C), and 106% of the average yield of individual strains (Fig. 4e).

### 4. Positive and negative pairwise interactions between members of the community

Emergent properties in microbial communities result from higher-order interactions and environmental conditions that drive community structure, function, assembly, stability, and evolution. However, the analysis of lower-order (pairwise) interactions can be useful to explain these complex processes (41, 66–68). To gain insights into the interactions driving the emergent properties of the Xilonen community, we evaluated pairwise interactions.

Negative interactions were addressed through an antagonism assay using the spot-on-lawn method (Fig. 5a). From the six possible pairwise combinations we found four antagonistic interactions. Strains A and B antagonized each other through weak and strong interactions, respectively; strains B and C antagonized mutually through weak interactions (Fig. 5a and 5c).

For the evaluation of positive interactions, we assessed pairwise cross-feeding. Bacterial growth was measured for each strain in cell-free spent medium from the remaining strains, compared to growth in self-spent medium and fresh LB. All strains grew in fresh LB, with an OD_600_ ranging from 1.54 to 1.96 (Fig. 5b). When grown in self-spent medium, only strain A exhibited growth (OD_600_ = 0.21). When cultivated in spent medium of different strains, we found four positive cross-feeding interactions. Metabolites produced by strains B and C supported the growth of strain A (OD_600_ = 0.33 and 0.85, respectively). Additionally, the spent medium of strain C facilitated the growth of strain B (OD_600_ = 0.25), and the spent medium of strain A supported the growth of strain C (OD_600_ = 0.52) (Fig. 5b and 5c).

### 5. Pairwise induction of biofilm formation

Complex architecture in bacterial colonies is mediated by molecules that also participate in biofilm formation (58, 59). For this reason, we evaluated pairwise induction of biofilm formation as a more direct approach to address this emergent property of the Xilonen community. Biofilm formation was determined qualitatively by visually detecting pellicle biofilms in the liquid-air interface, and quantitatively using the crystal violet method which targets surface-adhered biofilms (adhesion to microplate wells) (69). First, we compared the biofilm formation of individual strains, pairwise co-cultures, and the three-member community. Notably, only strain A formed a pellicle in single strain culture (Fig. 5a). When biofilm surface-adhesion was quantified, individual strains A, B, and C, exhibited 0.9, 0.13, and 0.06 mean OD_600_, respectively (Fig. 6b). Interestingly, we found that biofilm adhesion was not increased in pairwise co-cultures (mean OD_600_ of 0.13, 0.04, and 0.09 for AB, AC, and BC, respectively). Moreover, the three-strain ABC co-culture also failed to form a robust pellicle biofilm in standing liquid culture (mean OD_600_ = 0.07) (Fig. 6b). These findings suggest that the synthesis of compounds related to biofilm formation by the Xilonen community is contingent upon the structure provided by the agar medium.

We also evaluated pairwise induction of biofilm formation by cultivating single strains in conditioned medium (fresh LB amended with cell-free spent medium from the remaining strains at a 1:1 ratio), compared to fresh LB alone. Strain A developed a pellicle across all conditions (Fig. 6c). For strain B, a pellicle was developed in the presence of conditioned medium from Strain C (Fig. 6c and 6e), and bacterial growth was decreased in conditioned medium from strain A and B (self), compared to fresh LB (Fig. 6c). Strain C only developed a pellicle in conditioned medium of strain B (Fig. 6c and 6e). When biofilm surface-adhesion was quantified, strain A presented high biofilm adhesion in all conditions (mean OD_600_ from 1.01 to 1.86), showing a slight increase in the presence of conditioned medium from strain C compared to fresh LB (mean OD_600_ of 1.86 and 1.14, respectively) (Fig. 6d and 6e). For strains B and C, biofilm adhesion was not induced by any conditioned medium (Fig. 6d).

## Discussion

In this study, we present the assembly of a three-member higher-order synthetic community (SynCom) with emergent complex colony architecture, named the Xilonen SynCom as it comprises seed-endophytic strains from Mexican maize landraces. Our previous work showed that a fraction of the seed-endophytic bacteriome could contribute to maize fertility in traditional *milpa* agroecosystems, particularly during early stages of development (16–18). Plant fertility (i.e., the ability of plants to grow healthy and productive) results from the interplay of plant genotypes, environmental factors and complex interactions with and within associated microbial communities (46). In addition, cultural beliefs and traditions from rural regions in Mexico that use *milpa* systems strongly influence the selection of seeds and agricultural practices, unintentionally shaping plant biodiversity and the plant microbiome (17, 70). Understanding the contribution of the microbiota to plant health necessitates a holistic approach beyond the study of individual strains (39, 41). The strategy employed to obtain the Xilonen SynCom will serve as a generalizable approach for studying bacterial interactions in community assembly and for obtaining consortia for various applications. The Xilonen SynCom itself represents a promising model system to study the role of bacterial interactions in community assembly and its beneficial functions on native maize fertility.

Since SynComs are promising experimental tools in microbial ecology (42, 71) and their design is one of their main limitations (39–41), we developed a protocol for the bottom-up assembly of SynComs which is based on the detection of emergent collective properties. Many strategies have been used for selecting and combining strains in bottom-up approaches (42, 43, 72) where the assessment of emergent properties in a SynCom often comes last (37, 57, 73). However, here we show that detection of emergent traits in bacterial co-cultures could be an efficient criterion for SynCom assembly. Bottom-up approaches have also been combined with microfluidics and droplet encapsulation assays for the high-throughput screening of interactions in SynComs, resulting in a better understanding of the properties that drive community functions (47, 48). However, these approaches use highly specialized and often inaccessible equipment, including additional efforts for obtaining and processing a large number of samples and large datasets of microbial interactions and their outcomes (49, 50, 74). In our approach, screening ≈2% of all possible combinations was sufficient for detecting two communities with emergent collective functions. We recognize that our approach based on choosing interacting pairs for preparation of communities with additional members may limit our ability to identify more complex SynComs with emergent properties resulting from interactions among a larger group of strains. We also propose that our strategy could be optimized through the automation of bacterial mixture preparation, imaging, detection of emergent functions and phenotype comparisons between communities, individual members and subgroups.

Colony architecture in the Xilonen community depends on the coexistence of its three members. *In vitro* colony architecture is considered a proxy for biofilm formation in model bacterial species (75–77) since molecules shaping these macroscopic structures such as exopolysaccharides (EPS), extracellular proteins, and surfactants are components of the extracellular matrix (ECM) of biofilms found in nature (78). A structured biofilm generates heterogeneous microenvironments where physical and biological factors influence the fitness of strains and shape the community function, including its macroscopic properties (55, 79). Colony architecture in the Xilonen community was not accompanied by increased fitness of individual members. Hence, the emergent complex colony architecture is likely related to interactions resulting in the production of molecules that shape a structured ECM. Pairwise interactions (positive, negative, and biofilm induction) found between members of the Syncom may contribute to assembly, dynamics and stability, and could influence other biofilm properties such as spatial distribution of the populations inside the colony biofilm (80). Microenvironments created inside biofilms due to the physical arrangement of cells could also impact cell differentiation (81–83). Thus, ecological traits such as division of labor could be promoted in the Xilonen community. Future studies will address the spatio-temporal dynamics of cells within the biofilm, along with metabolomics analyses to depict the molecular mechanisms driving the presence of a complex colony architecture in the SynCom.

After decades of research on plant growth-promoting bacteria (PGPB) and the development of microbial-based products to support plant health (25, 26), the effectiveness of these products in the field remains largely uncertain (21–24). Inoculants have recently incorporated microbial consortia to improve outcomes, but formulations lack ecological validation (20, 25, 26). Additionally, the study of microbial interactions and their impact on community functions are generally neglected. Instead, it is assumed that mixing ‘biocompatible’ PGPB strains (i.e., strains lacking antagonistic interactions in the spot-on-lawn method), or strains presenting complementary functions will ultimately result in additive or synergistic traits (27–36). However, community functions cannot be predicted as the sum of its parts (84) and, moreover, negative interactions are common in community assembly and stability (66, 85). Indeed, we show that strains selected in the Xilonen community display negative interactions that do not affect community stability, since stability is an emergent property that arises from the intrinsic dynamics of microbial systems (86–88). Introduced microbes in agriculture should be able to perform their functions in varying environmental conditions and in the presence of native soil microbiota, therefore, formulating inoculants through synthetic ecology approaches is a promising solution to develop ecologically-relevant and functionally robust microbial products.

## Conclusion

Our study introduces a novel three-strain synthetic community (SynCom) named “Xilonen”. By tapping into the richness of seed-endophytic bacteria from maize landraces, we constructed SynComs that mirror complex functions observed in natural communities. Our systematic bottom-up approach prioritizes the selection of strains based on anticipated properties of the assembled higher-order community, which could translate to enhanced performance in agricultural settings. Notably, our focus on biofilm formation, a key factor in root colonization, underscores the significance of these SynComs in agricultural applications. The Xilonen community exemplifies the potential of combinatorial screenings to identify communities with emergent properties aligned with desired outcomes. These SynComs represent invaluable model systems for advancing our understanding of how microbial communities influence maize development, paving the way for innovative agricultural practices and sustainable crop management strategies.

## Methods

### Bacterial strains and culture conditions

Strains used in this study are shown in Supplementary Table S1. For all experiments, strains were streaked from cryostocks on LB agar (10 g L^-1^ tryptone, CRITERION™, 5 g L^-1^ yeast extract, Sigma-Aldrich and 5 g L^-1^ NaCl) and incubated for 2 days at 30 °C. For slow-growing strains (NME36, NME37, NME100, NME117, NME186, NME233 and NME235), one colony was picked into 50 mL of LB liquid medium and grown at 30 °C and 200 rpm for 40-42 h. For the rest of the strains, one colony was picked into 20 mL of LB liquid medium and grown at 30 °C and 200 rpm for 16-18 h. Rifampicin 50 μg/mL (rif50), tetracycline 10 μg/mL (tet10), or chloramphenicol 10 μg/mL (cam10) were added when needed.

### Combinatorial screening of emergent colony architecture in bacterial communities

The general strategy for the bottom-up assembly of SynComs with emergent properties is shown in Figure 1a. For the preparation of bacterial strain combinations, 10 mL of liquid cultures from all strains (Table S1) were centrifuged (4000 rpm for 10 min), washed, and suspended in 10 mL of sterile PBS. Then, the Optical Density at 600 nm (OD_600_) was adjusted to 1 ± 0.1. These suspensions were mixed into combinations in microtiter 96-well plates (AXYGEN, No. P-DW-11-C-S) at an equal volume ratio with a final volume of 350 μL. Once all combinations were prepared, plates were covered with a sealing mat (AXYGEN, No. AM-2ML-RD), vortexed to ensure a homogeneous mixture, and spun briefly before re-opening. Then, 5 μL of each of the 96 wells was spotted on a previously air-dried LB plate (15 cm diameter). Inoculated spots were air-dried for 15 min, incubated at 30 °C for 3 days, and imaged using a digital camera and a copy stand. Mixed colonies were visually screened for emergent colony architecture. Emergent colony architecture was determined when the morphology of a mixed colony differed from the morphology of all individual members (Fig. 1b). Emergent architecture of selected colonies was confirmed on separate LB plates to avoid the effect of metabolite diffusion through the agar, and communities with confirmed emergent colony morphology were used for a subsequent step of combination prep and screening (community *k* = n+1, Figure 1a and 1c). All spot-inoculations were performed in triplicate.

### Genome sequences and species-level identification

DNA from *Bacillus* sp. NME155, *Burkholderia* sp. XM7, and *Pseudomonas* sp. GW6 was isolated from 2 mL of overnight liquid cultures grown in LB medium at 30 °C. Cells were harvested by centrifugation (8000 rpm, 3 min), and DNA was isolated from the pellet using the Power Soil Pro Kit (QIAGEN). The first step of the protocol was modified for our samples: pellets were suspended in 800 μL of solution CD1 and transferred to the PowerBead Pro Tube. Then, the protocol was followed as described in the manufacturer’s handbook. Pair-end genome sequences (2 x 150 bp) were obtained using the Illumina Miniseq platform (Laboratorio de Genómica Microbiana, CIAD Mazatlán, Sinaloa, Mexico). Illumina reads were adapter and quality trimmed using FastQC v0.11.2. After trimming, the quality was assessed one more time using FastQC v0.11.2. Genomes were assembled using SPAdes v3.15.2 (89), and the quality of each assembly was verified using QUAST v5.0.2 and CheckM2 v.0.1.2 (90, 91).Species-level identification of each strain was made using the Microbial Genomes Atlas (MiGA) web server v1.2.18.2 (92, 93), and digital DNA–DNA hybridization (dDDH) was calculated using the Type (Strain) Genome Server (TYGS) online platform (94). Before GenBank submission, contigs < 200 bp were eliminated from the genome assemblies.

### Spontaneous rifampicin-resistant variant of *Bacillus pumilus* NME155

To recover Bp_NME155 from the Xilonen community, we generated a spontaneous rifampicin-resistant variant as described in (16). Briefly, cells from 40 mL of an overnight liquid culture were harvested by centrifugation (8000 rpm and 4 °C for 10 min). The resulting pellet was suspended in 2 mL of fresh LB medium, and 500 μL of the suspension was plated on LB medium with rifampicin (50 μg/mL). Plates were air-dried and incubated in darkness at 30 °C for 2 days. Next, 5 rif^R^ colonies were isolated and a variant with a similar growth rate (LB liquid medium) and colony morphology, compared to the wild-type strain, was selected. Additionally, the presence of emergent colony architecture in the Xilonen community assembled with the selected rif^R^ variant was verified. The strain was grown in LB medium and preserved with 15% glycerol at 80 °C.

### CFU counts from colony biofilms

Colony biofilms of *Bacillus pumilus* NME155 (Bp_NME155), *Burkholderia contaminans* XM7 (Bc_XM7), and *Pseudomonas* sp. GW6 (P_GW6), were grown as described above. For inoculation of the community, cultures were mixed on a 1:1:1 volume ratio and 5 μL of the mixture was used for spot inoculation. CFU counts at day 0 were estimated by plating 10-fold serial dilutions of the liquid cultures used for spot-inoculation. For CFU counts from colony biofilms, three colonies of individual strains and the community were cut off the agar plate with a sterile scalpel and transferred to a culture tube (25 mm x 150 mm) with 5 mL of sterile PBS. Colonies were suspended by vortexing twice for 1 min, and 10-fold serial dilutions of each suspension were plated on LB, LB rif50, and LB tet10. Plates were incubated at 30 °C and CFU counts were assessed after 1 to 2 days. In the mixed colonies, CFUs of strain A (Bp_NME155), strain B (Bc_XM7), and strain C (P_GW6) were estimated by CFU counts on LB rif50, LB tet10, and LB, respectively.

### Antagonistic interactions assay

Antagonistic interactions were assessed using the spot-on-lawn assay (95). Briefly, overnight liquid cultures of strains *Bacillus pumilus* NME155 (Bp_NME155), *Burkholderia contaminans* XM7 (Bc_XM7) and *Pseudomonas* sp. GW6 (P_GW6) were washed as described above, and OD_600_ was adjusted to 1 ± 0.1. The sensitive strain was inoculated as a lawn mixed at 1:1000 volume in LB agar before pouring into Petri dishes. Plates inoculated with each “lawn-strain” were air-dried for at least 30 min, and 5 μL of each “spot-strain” were inoculated on top. Plates were air-dried for another 15 min and incubated at 30 °C. Antagonistic interactions were evaluated after 2 days of incubation and they were classified as strong antagonisms (defined halo) and weak antagonisms (dim halo). Each interaction was performed in triplicate.

### Cross-feeding assay

Pairwise cross-feeding was evaluated by measuring the growth of each strain in cell-free spent medium prepared from a different strain. For preparation of cell-free spent medium, one colony (strain A: Bp_NME155, strain B: Bc_XM7 and strain C: P_GW6) was picked into 100 mL of LB medium and grown at 30 °C and 200 rpm for 5 days. Next, cells were removed by centrifugation (10,000 rpm for 20 min, twice), and the supernatant was collected and filter-sterilized using a 0.2 μm pore size. To verify sterilization, 10 μL of each spent medium was plated on fresh LB and incubated at 30 °C for 5 days. For inoculation, overnight liquid cultures of strains Bp_NME155, Bc_XM7 and P_GW6 were grown and washed as described in the previous sections. OD_600_ was adjusted to 1 ± 0.1 and 3 μL were used to inoculate 3 mL of fresh LB or spent medium of strains A, B and C. Cultures were grown at 30 °C and 200 rpm, and bacterial growth in each medium was assessed by measuring OD_600_ after 2 days. All cultures were performed with three replicates.

### Biofilm formation assays

Pairwise induction of biofilm formation was assessed in two separate assays, 1) in bacterial co-cultures (*k* = 2, 3) in fresh LB medium compared to single strain cultures, and 2) in single strain cultures grown in conditioned medium (2X LB medium mixed with spent medium from a different strain culture in a 1:1 ratio). Biofilm formation in both assays was quantified using the crystal violet method with modifications (16, 69). Briefly, overnight cultures of strains Bp_NME155, Bc_XM7, and P_GW6 were grown, washed, and adjusted to OD_600_ 1 ± 0.1. For assessing biofilm formation in bacterial co-cultures, each single strain, pairwise co-culture, and the community (mixed at an equal volume ratio) were diluted 1:100 in fresh LB medium. For the experiment using conditioned medium, each strain was diluted 1:100 in fresh LB medium or conditioned medium prepared with spent medium from each of the individual strains. Next, 100 μL of inoculated media were transferred into sterile non-tissue culture treated 96-well microtiter plates, and inoculated plates were incubated at 30 °C without agitation for 3 days. Planktonic cells were removed, plates were washed as described in (69), and biofilms were stained with 250 μL of 0.1% (w/v) crystal violet for 10 min at room temperature. After incubation, plates were washed twice and stained biofilms were dissolved with 250 μL of 30 % (v/v) glacial acetic acid for 15 min. For quantification, 125 μL of the dissolved and stained biofilm was transferred to a clear flat-bottom microtiter plate, and OD_600_ was measured using a plate reader (Varioskan Lux, Thermo Scientific). For both assays, representative microtiter wells were imaged using an Axio Zoom.V16 stereo microscope with an incorporated Axiocam 105 color 429 (Carl Zeiss Microscopy, Oberkochen, Germany) before quantifying biofilm formation. Wells with non-inoculated medium were used as control. At least 3 replicates were used on each assay.

## Acknowledgements

This project was financed by CONAHCYT Ciencia de Frontera project No. 39589, granted to GOA and JR. GG received a fellowship from CONAHCYT (761832). Authors thank Xcaret Molina and Martha Guerrero for technical assistance.

