## Supplementary Material for "Harnessing Emergent Properties of Microbial Consortia: Assembly of the Xilonen SynCom"

### Supplementary tables

**Supplementary Table S1. Bacterial strains used for the combinatorial screening of interactions.**

| Code | Strain | Reference |
|---|---|---|
| NME9 | *Bacillus* sp. | [1] |
| NME26 | *Bacillus* sp. | |
| NME32 | *Domibacillus* sp. | |
| NME36 | *Bacillus* sp. | |
| NME37 | *Domibacillus* sp. | |
| NME52 | *Bacillus* sp. | |
| NME63 | *Bacillus* sp. | |
| NME85 | *Bacillus* sp. | |
| NME100 | *Paenibacillus* sp. | |
| NME101 | *Bacillus* sp. | |
| NME117 | *Bacillus* sp. | |
| NME135 | *Peribacillus* sp. | |
| NME155 | *Bacillus* sp. | |
| NME186 | *Paenibacillus* sp. | |
| NME233 | *Bacillus* sp. | |
| NME235 | *Paenibacillus* sp. | |
| NME239 | *Paenibacillus* sp. | |
| NME246 | *Bacillus* sp. | |
| NME247 | *Peribacillus* sp. | |
| XM5 | *Burkholderia* sp. | [2] |
| XM7 | *Burkholderia* sp. | |
| XM13 | *Burkholderia* sp. | |
| GW1 | *Pseudomonas* sp. | [3] |
| GW6 | *Pseudomonas* sp. | |
| GW9 | *Pseudomonas* sp. | |
| GW12 | *Pseudomonas* sp. | |

**Supplementary Table S2. Genomic sequencing, assembly, and annotation statistics of members of the SynCom**

| | **Species** | ***Bacillus pumilus* NME155** | ***Bukholderia contaminans* XM7** | ***Pseudomonas* sp. GW6** |
|---|---|---|---|---|
| **Sequencing** | **Total reads** | 1,246,602 | 3,342,712 | 1,456,018 |
| | **Mean reads length (bases)** | 93 | 85 | 102 |
| | **Coverage (×)*** | 31 | 56.04 | 27 |
| **Assembly** | **Assembled genome size (bp)** | 3,796,551 | 5,069,692 | 5,488,062 |
| | **Number of contigs** | 160 | 8,997 | 574 |
| | **N50 value** | 97,169 | 1,117 | 25,285 |
| | **L50 value** | 12 | 644 | 66 |
| | **GC content (%)** | 41.59 | 61.63 | 62.5 |
| | **Completeness (%)** | 100 | 66.39 | 100 |
| | **Contamination (%)** | 0.03 | 7.48 | 0.38 |
| **Identification** | **Closest type strain (NCBI RefSeq assembly no.)** | GCF_900186955.1 | GCF_000987075.1 | GCF_002741105.1 |
| | **Average Nucleotide Identity ANI (%)** | 95.36 | 99.92 | 91.89 |
| | **digital DNA-DNA Hybridization dDDH (%)** | 63.9 | 98.9 | 44.8 |
| **Annotation** | **Number of coding sequences** | 4,056 | ** | 5,489 |
| | **Number of RNAs** | 86 | ** | 79 |
| | **Annotated gene number** | 1,385 | ** | 1,591 |

Sequencing metrics shown here resulted after filtering and clean-up steps.
*, sequencing coverage was calculated as (total reads*mean read length)/assembled genome size
**, genome assembly was too low quality for annotation with RAST

**Supplementary figures**

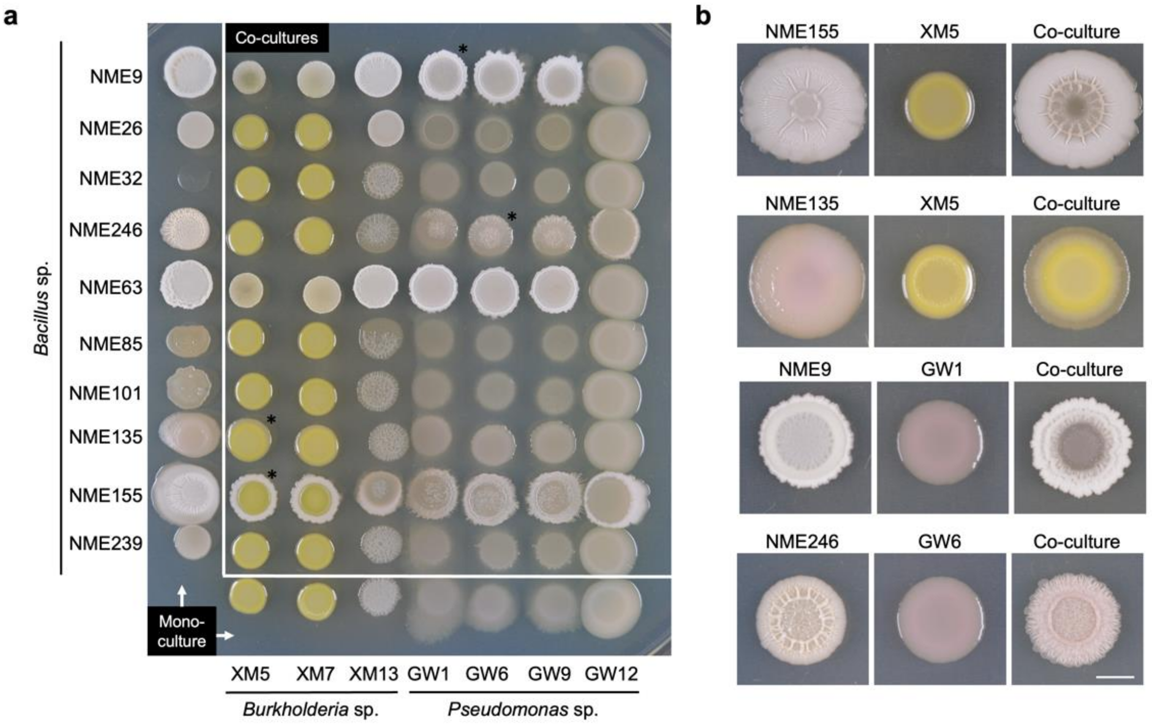


**Supplementary Figure S1. Screening of communities *k* = 2. a,** Representative plate for screening 70 co-cultures *k* = 2 after 3 days of incubation. Colonies with asterisks are shown in part b. **b,** Representative communities *k* = 2 with emergent colony morphology, selected for a subsequent step of combination prep and screening. Colonies were grown in separate LB plates and pictures were taken after 3 days of incubation at 30 ºC. Scale bar: 5 mm.

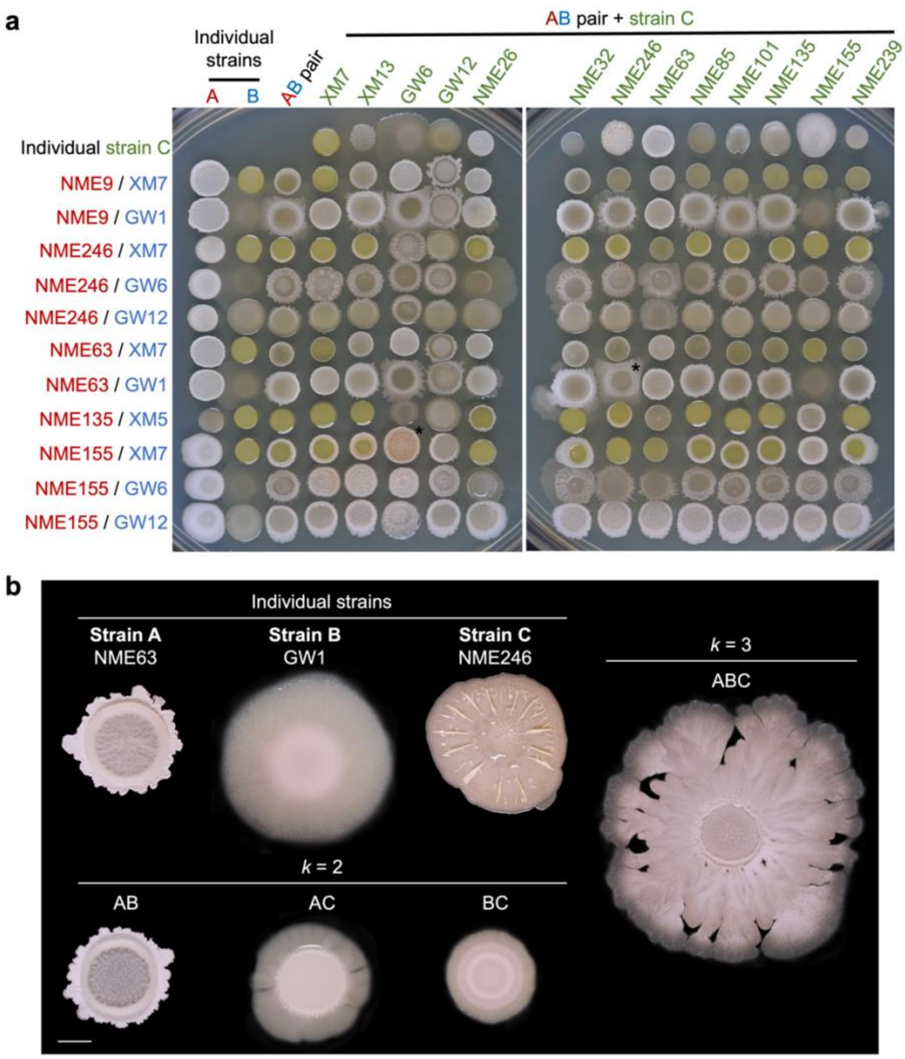


**Supplementary Figure S2. Screening of communities *k* = 3. a,** Representative plates for screening 143 co-cultures *k* = 3 after 3 days of incubation. The phenotype of colonies with asterisks was confirmed in separate plates. **b,** Community *k* = 3 presenting emergent colony spreading. Colonies were grown in separate LB plates and pictures were taken after 3 days of incubation at 30 ºC. Scale bar: 5 mm.

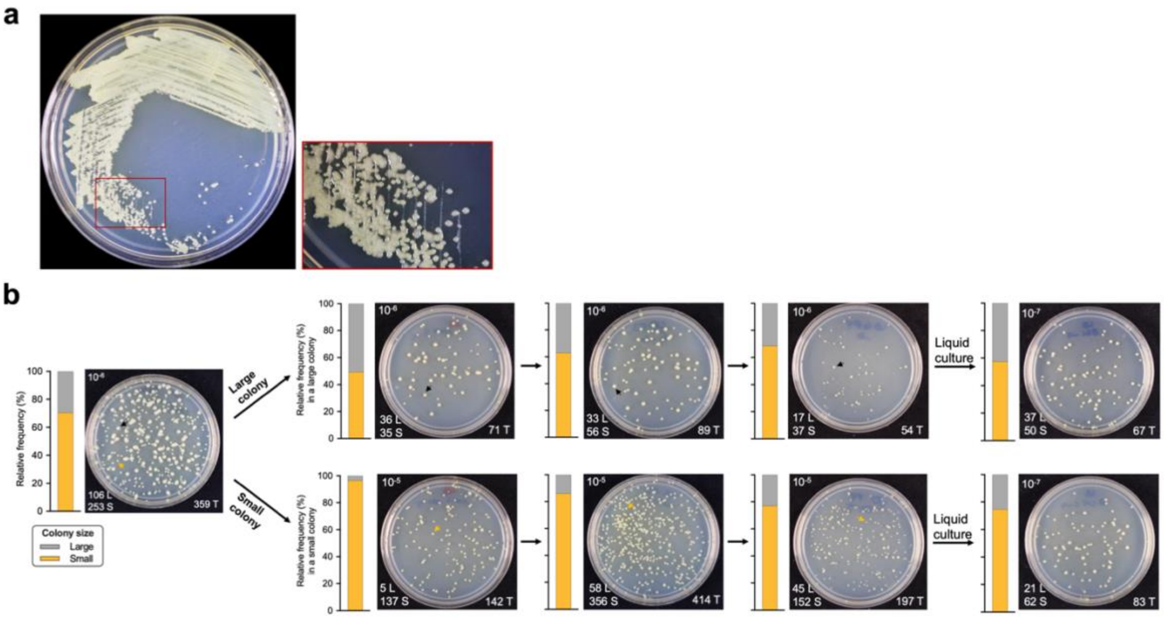


**Supplementary Figure S3. The bacterial culture of strain *Burkholderia contaminans* XM7 presents 2 morphologies. a,** Strain Bc_XM7 streaked on LB media presenting two colony morphologies. **b**, Proportion of large and small colonies in bacterial cultures of Bc_XM7. Serial dilutions 1:10 of an overnight liquid culture of Bc_XM7 were plated on LB media and the proportion of large and small colonies were calculated. Then, one large and one small colony were suspended in 200 µL of sterile PBS, serial dilutions 1:10 were plated, and the proportions of both colony morphologies were calculated. This purification step was repeated three times. After the third purification step, colonies were picked into liquid media and grown overnight; serial dilutions were prepared form these liquid cultures for calculation of proportions. CFU counts of large (L), small (S), and total (T) colonies are shown in each picture. All pictures were taken after 2 days of incubation at 30 ºC.

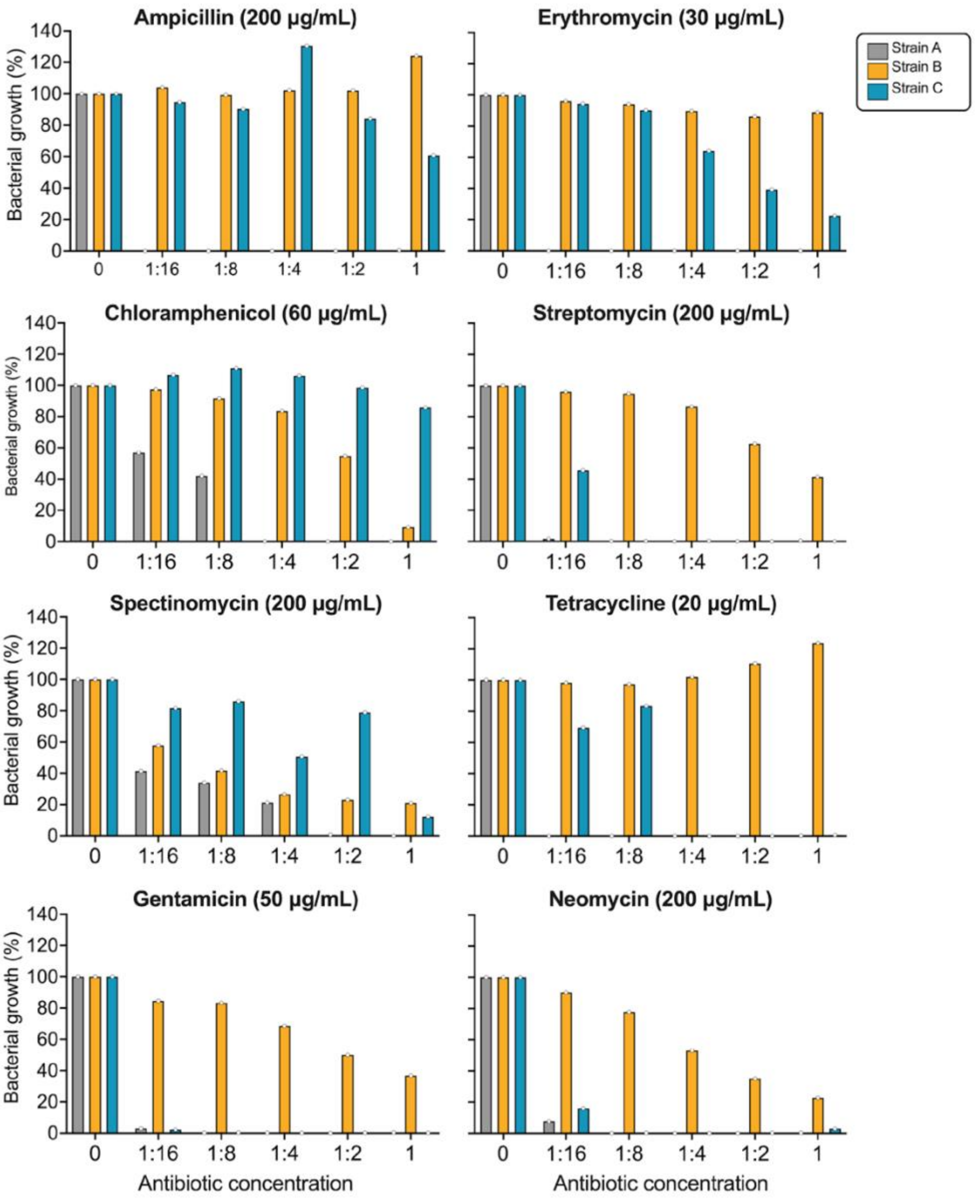


**Supplementary Figure S4. Screening of antibiotic resistance for individual strains in the SynCom.** Minimum inhibitory concentration assays were performed using 8 antibiotics and three strains. The highest concentration (identified as ‘1’ on the X-axis) of each antibiotic is shown in parentheses. Strains were inoculated in 250 µl of fresh LB at a dilution factor of 1:500 from overnight liquid cultures previously washed with PBS. Microtiter plates were incubated without agitation at 25 ºC and Optical Density at 600 nm ($OD_{600}$) was measured after 3 days. Bacterial growth ($OD_{600}$) in antibiotics is shown as a percentage (%) of growth in LB media without antibiotics. Strain A: *Bacillus pumilus* NME155 (Bp_NME155); Strain B: *Burkholderia contaminans* XM7 (Bc_XM7) and Strain C: *Pseudomonas* sp. GW6 (P_GW6).

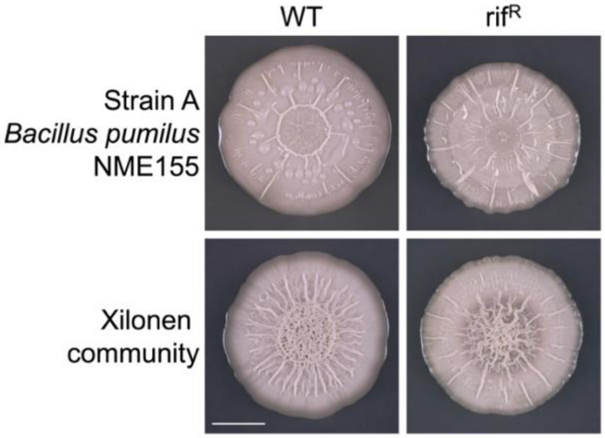


**Supplementary Figure S5. Colony architecture of the Xilonen community using *Bacillus pumilus* NME155 Wild Type (WT) or the rifampicin-resistant variant (rif$^R$).** Strains were grown in 20 mL of LB media at 30 ºC and 200 rpm for 16 h. Bacterial cultures were washed with PBS (see methods) and the optical density at 600 nm ($OD_{600}$) was adjusted to 1 ± 0.1. The community was assembled with either the WT or the rif$^R$ variant of strain *B. pumilus* NME_155 by mixing the bacterial cultures at a 1:1:1 volume ratio, and 5 µL of each strain and the community were spotted on independent LB agar plates. Pictures were taken after 3 days of incubation at 30 ºC. Scale: 0.5 cm.

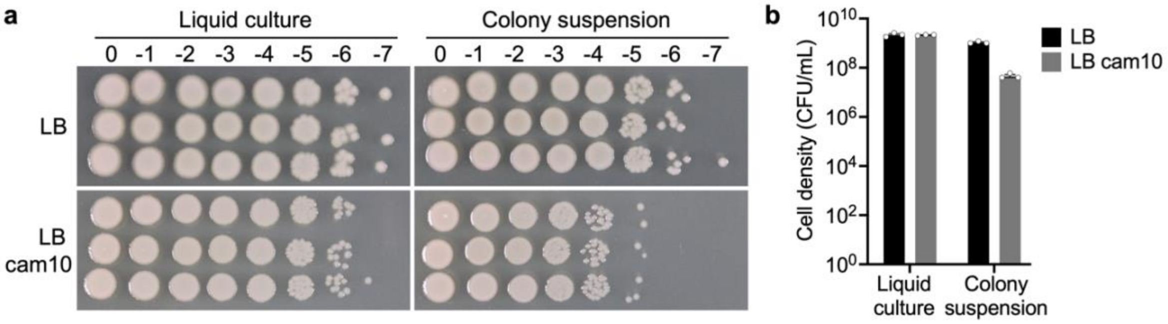


**Supplementary Figure S6. Chloramphenicol resistance of *Pseudomonas* sp. GW6 (strain C) is decreased in colony biofilm, compared to liquid culture. a,** CFU counts of *Pseudomonas* sp. GW6 were evaluated using 10-fold serial dilutions from triplicate liquid cultures and colony suspensions, which were spotted on LB and LB cam10. **b,** Cell density of liquid cultures and colony suspensions measured on LB and LB cam10. A washed overnight liquid culture was used for this experiment. For colony suspensions, a 3-day-old colony was used (5 µl spot inoculated in LB agar).
